# assayM: a web application to monitor mutations in COVID-19 diagnostic assays

**DOI:** 10.1101/2020.12.18.423467

**Authors:** Raeece Naeem, Arnab Pain

## Abstract

**Summary:** Reverse Transcriptase – Polymerase Chain Reaction (RT-PCR) is the gold standard as diagnostic assays for the detection of COVID-19 and the specificity and sensitivity of these assays depend on the complementarity of the RT-PCR primers to the genome of the severe acute respiratory syndrome coronavirus 2 (SARS-CoV-2). Since the virus mutates over time during replication cycles, there is an urgent need to continuously monitor the virus genome for appearances of mutations and mismatches in the PCR primers used in these assays. Here we present assayM, a web application to explore and monitor mutations introduced in the primer and probe sequences published by the World Health Organisation (WHO) or in any custom-designed assay primers for SARS-CoV-2 detection assays in globally available SARS-CoV-2 genome datasets.

**Availability and implementation:** assayM is available on https://grafnet.kaust.edu.sa/assayM as a web application and also as an open-source R shiny application, downloadable from https://github.com/raeece/assayM

## 1 Introduction

The gold standard for diagnosing the SARS-CoV2 virus is the RT-PCR. The PCR method involves a forward primer, reverse primer and probe to detect the presence of SARS-CoV2 virus infection. There are a number of PCR assays published in the literature (Khan and Cheung, 2020),(Wang, et al., 2020) and recommended by the World Health Organisation (WHO) (https://www.who.int/docs/defaultsource/coronaviruse/whoinhouseassays.pdf). The SARS-CoV2 virus continues to mutate and mutations appear in the regions where the assay primers bind to the RNA strand of the virus. As mutations accumulate in the target regions of the PCR primer and the probe binding region in the virus genome, the diagnostic efficacy of these assays gets reduced leading to false negative results in these tests(Wang, et al., 2020). It is imperative to monitor the mutations acquired in the target region to identify and redesign the primers to main a high level of sensitivity and specificity of the diagnostic assays. Samples of SARS-CoV2 virus strains are being sequenced across the world every day and made public through Genbank(Sayers, et al., 2020) and GISAID(Shu and McCauley, 2017). In order to track these mutations, we developed an exploratory web application assayM to visualize the impact of these mutations in the published RT-PCR primers and probes. This web application would be useful in designing PCR Primers with the least number of mutations. This application eases the time-consuming process of downloading newly available sequences and evaluating the diagnostic assays.

The web application is built using R language and scripts to compile the database of mutations is written in python and available to anyone to customize them according to their needs.

## 2 assayM web application

The core components required of the assayM web application are

Compiled list of published PCR primers and probe sequences.

Catalog of curated Single Nucleotide Polymorphisms (SNPs) and Insertions and Deletions (Indels.)

Metadata such as date of collection and geographical origin of sequences.

An integrated database and a visualization tool to search and filter regions of interest and track their changes over time.

PCR primers and probes were collated from (Khan and Cheung, 2020) where they evaluated 27 diagnostic probes and primers using a snapshot of 17,000 viral sequences available at that time. (Wang, et al., 2020) also evaluated published assays based on 31,421 sequences available in July, 2020. As of writing today there are in excess of 270,000 viral sequences available in the global repositories and continues to grow in thousands every two days. A catalog of curated SNPs and Indels are compiled from sequences available on Genbank and GISAID and good quality variants are made available for analysis every day on 2019 Novel Coronavirus Resource (Zhao, et al., 2020). The assayM python scripts pulls the curated multi-sample VCF file and its associated metadata from 2019 Novel Coronavirus resource on a weekly basis and builds an SQL database for querying and visualisation of the regions where the mutations occur.

### 2.1 Exploring mutations in primer regions

The core functionality of assayM is to visualize the primer and probe regions and summarise the number of strains that have mutations in the target region. A genome browser with gene annotations is shown along with the primer and probe regions. (Fig 1) The user has the ability to input and search their custom primer and probe sequences for a match against the SARS-CoV2 Wuhan-Hu-1 reference sequence (NCBI accession NC_045512). The user can view any region of interest for mutations and obtain the mutation statistics. The bar graphs summarise the proportion of sequences that contain mutations against the total number of sequences deposited in the sequence repositories every month. By visualizing the mutation counts and proportion one can choose the primer regions when designing new primers and discard existing primers if there is an indication of too many mutations that may result in reduction of specificity and sensitivity of the assays.

**Fig. 1.**
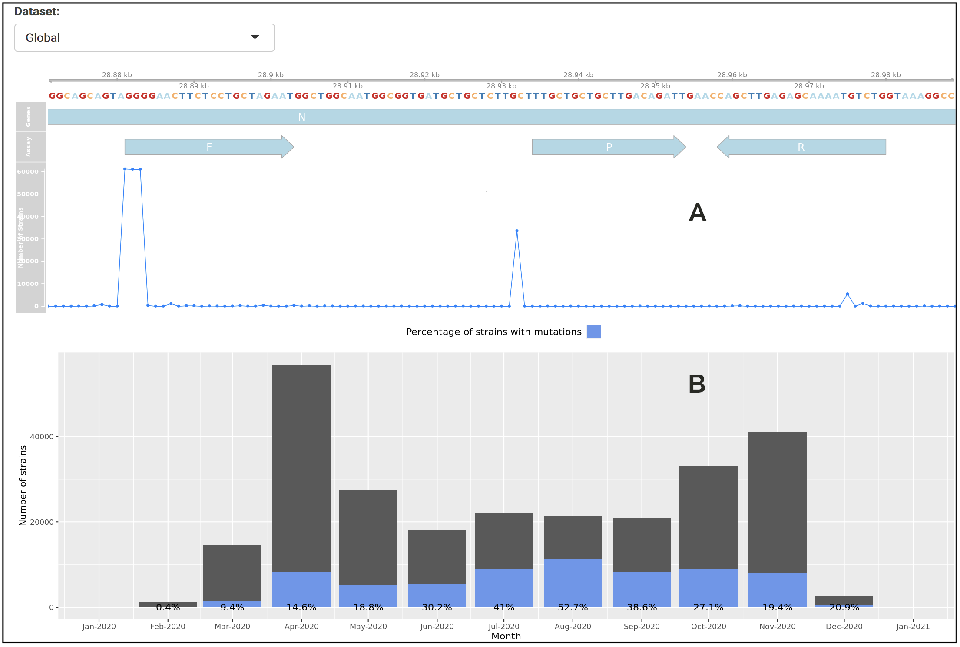
Explore mutations in primer regions. A) This example show the number of strains with mutations in the CN-CDC-N primer set. B) Shows the proportion of strains that had mutations every month against the total number of strains deposited.

### 2.2 Compilation of published assays

A total of 34 diagnostic assays were compiled from literature review (Khan and Cheung, 2020; Wang, et al., 2020) and assays published on WHO website. Assays are listed in a table format where one can sort the primer sets according to most or least number of mutations. Table 1 highlights 6 primer set where three of them contains the greatest number of mutations and the other three with the least number of mutations among the compiled set of 34 assays.

**Table 1.** Table of assays with most number to least number of mutant strains.

| \|S\| | Assay | Mutant strains | Assay Country | Gene Target |
| --- | --- | --- | --- | --- |
| 1 | CN-CDC-N | 67871 | China | N |
| 2 | Young-S | 7135 | Singapore | S |
| 3 | SC2 | 6541 | US | N |
| 1 | Won-ORF1ab | 608 | South Korea | ORF1ab |
| 2 | CN-CDC-ORF1ab | 903 | China | ORF1ab |
| 3 | Pasteur-ORF1ab-1 | 909 | France | ORF1ab |

Total number of strains deposited at the time of compiling this table is 261,988

### 2.3 Detailed view on SNPs and Indels

A separate tab of the application lists all the mutations occurring across all regions of the SARS-CoV2 virus genome. For each mutation the genomic co-ordinates, strain name, sample collection date, country of origin, mutation type (SNP/Indel), reference and alternate alleles are displayed. The user can filter the table by genomic co-ordinates, country, collection date or mutation type.

### 2.4 Conclusion

Through assayM application we have included the most essential statistics on mutations related to diagnostic RT-PCR assays during the chaging landscape of the COVID-19 pandemic. As the source code is written in R shiny framework and the python scripts required to compile the mutation database is made available one can customize the R code to include more explorative charts and graphs on mutations in SARS-CoV2 virus.

## Acknowledgements

We thank KAUST Pathogen Genomics group members for their comments and sup-port.

## References

Khan, K.A. and Cheung, P. Presence of mismatches between diagnostic PCR assays and coronavirus SARS-CoV-2 genome. Roy Soc Open Sci 2020;7(6).

Sayers, E.W., et al. GenBank. Nucleic Acids Res 2020;48(D1):D84–D86.

Shu, Y.L. and McCauley, J. GISAID: Global initiative on sharing all influenza data - from vision to reality. Eurosurveillance 2017;22(13):2–4.

Wang, R., et al. Mutations on COVID-19 diagnostic targets. Genomics 2020.

Wang, X.L., et al. Limits of Detection of 6 Approved RT-PCR Kits for the Novel SARS-Coronavirus-2 (SARS-CoV-2). Clin Chem 2020;66(7):977–979.

Zhao, W.M., et al. The 2019 novel coronavirus resource. Yi Chuan 2020;42(2):212–221.

